# Data coverage and model formulation reshape quantitative interpretations of bacterial transcriptional regulation

**DOI:** 10.64898/2026.08.31.748186

**Authors:** Syue-Ting Antony Kuo, Chao-Ping Hsu, Hsin-Hung David Chou

## Abstract

Thermodynamic models quantitatively describe interactions between transcription machinery and bacterial promoters. Contrary to conventional understanding, model analysis by Parisutham et al. (2025) attributes transcriptional inhibition by repressors to overstabilization of the RNA polymerase–promoter complex rather than prevention of its formation. Moreover, it suggests an inverse scaling relationship between basal promoter strength and transcriptional fold change, applicable to both repressor- and activator-mediated regulation. To reevaluate findings from this study, we systematically analyze empirical data and compare its framework with conventional thermodynamic models. In contrast to the inverse scaling relationship, data across multiple sources exhibit a peaked tradeoff between basal promoter strength and fold change, underscoring the importance of broad data coverage in revealing the full pattern required for reliable model inference. Furthermore, we identify the model assumption responsible for the apparent inverse scaling and misinterpretation of regulatory mechanisms. Relaxing this assumption enables the model to capture the peaked tradeoff and yield inferences consistent with established mechanisms of transcriptional repression and activation. We further derive a mathematical solution that connects basal expression to fold change for both repressor- and activator-regulated promoters. Our results underscore the importance of broad data coverage to avoid a “blind men and the elephant” interpretation and establish basal promoter strength as a key design parameter governing transcriptional regulation.

## Introduction

Cells adjust their physiological responses to changing environments through gene regulation [1]. In bacteria, gene expression is regulated primarily at the transcription level through interactions among promoters, RNA polymerase (RNAP), and transcription factors (TFs) [2]. Transcription begins when a σ factor, mainly the housekeeping σ^70^, directs RNAP to bind the promoter region and unwind double-stranded DNA for RNA synthesis [3]. Core promoter elements, particularly the sequence composition of the −35 and −10 elements, determine the basal promoter strength, while TFs repress or activate transcription by modulating RNAP–promoter binding [2,4]. Transcriptional repressors (e.g., TetR) reduce gene expression by blocking RNAP access to promoters during transcription initiation or by impeding transcription elongation (**Fig 1A**) [2,5]. Transcriptional activators, on the other hand, facilitate formation of the transcription initiation complex through direct contact with RNAP (e.g., Crp, LuxR; **Fig 1B**) or by remodeling promoter DNA conformation (e.g., CueR) [6–8]. To quantitatively dissect the regulatory logic, modeling approaches have been developed that fall broadly into black-box or white-box strategies. Black-box models, such as deep learning, achieve high predictive accuracy but lack mechanistic interpretability [9,10]. In contrast, white-box models, exemplified by thermodynamic models, provide transparent frameworks for TF–RNAP interactions and promoter sequence–function relationships [11,12]. By explicitly parameterizing molecular interactions, thermodynamic models yield mechanistic insights in addition to predicting transcriptional output.

**Fig 1.**
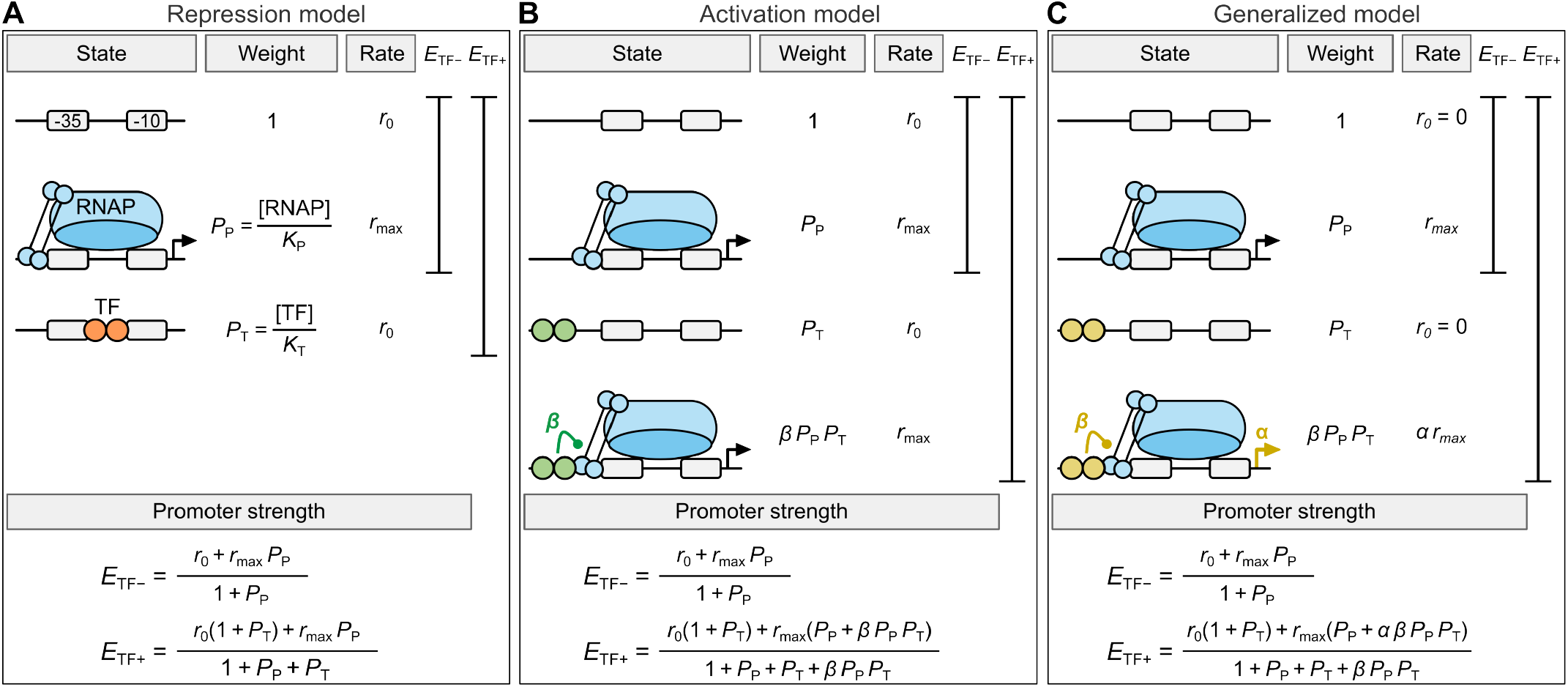
Thermodynamic models of transcriptional regulation. Schematics of the repression **(A)**, activation **(B)**, and generalized **(C)** models. Promoter states (from top to bottom: unbound, RNAP-bound, TF-bound, co-bound), their corresponding Boltzmann weights and transcription rates, the states contributing to basal (*E*_TF−_) and regulated (*E*_TF+_) expression, and the equations for computing *E*_TF−_ and *E*_TF+_ are indicated in each panel. The generalized model (GM) sets *r*_0_ = 0. The revised generalized models, GM* and GM**, are identical to GM except that GM* treats *r*_0_ as a free parameter and GM** further sets *α* = 1. Parameter definitions: *P*_P_ and *P*_T_, Boltzmann weights of RNAP-bound and TF-bound states, respectively; [RNAP] and [TF], concentrations of RNAP and of binding-competent TF, respectively; *K*_P_ and *K*_T_, dissociation constants of RNAP and TF, respectively; *r*_0_, rate of background transcription beyond the defined promoter region; *r*_max_, transcription rate of the RNAP-bound state (*r*_0_ ≪ *r*_max_); *α*, TF influence on promoter escape; *β*, TF influence on RNAP–promoter binding.

Thermodynamic modeling can be traced back to 1974, when von Hippel and colleagues applied it to infer protein–DNA binding mechanisms from *in vitro* binding and *in vivo* gene expression data [13,14]. Their analysis showed that *lac* operon regulation could be rationalized only when non-specific LacI–DNA binding was taken into account. Subsequently, Ackers et al. expanded this framework by incorporating cooperative protein–protein interactions in modeling the lytic–lysogenic switch of bacteriophage λ [15]. Their study demonstrated that *in vivo* gene expression could be predicted solely from binding energies measured *in vitro*. With the emergence of synthetic biology in the 2000s, Buchler et al. explored the potential of thermodynamic modeling in designing genetic circuits by computationally adjusting basal promoter strength and the relative positions of TF-binding sites (TFBSs) [16]. To generalize this approach, Bintu et al. proposed a single parameter, the regulation factor, to quantify the influence of repressor and activator TFs on gene expression [11,17]. To acquire large-scale promoter sequence– function data, Kinney et al. in 2010 developed sort-seq, a massively parallel reporter assay (MPRA) that combines mutational scanning, fluorescence-activated cell sorting, and deep sequencing [18]. Model fitting to their sort-seq data enabled quantification of RNAP–promoter and TF–TFBS binding energies at single-base resolution. Brewster et al. then used this information to engineer the *lac* promoter, demonstrating rational design of transcription activity across a wide dynamic range [19]. Since then, thermodynamic modeling and MPRAs have been applied jointly to unravel TF regulatory mechanisms, dissect promoter architecture and design principles, elucidate promoter sequence evolution, and construct synthetic promoters [20–30].

The conventional thermodynamic framework accounts for the effects of repressors and activators using the repression model (RM; **Fig 1A**) and the activation model (AM; **Fig 1B**), respectively [11,12]. As the two models assume different TF–RNAP interactions, modeling requires the subject TF to be classified *a priori* as an activator or a repressor. However, such classification is often infeasible for poorly characterized TFs. To circumvent this, Brewster and colleagues developed a generalized model (GM) applicable to both repressors and activators, formulated to infer TF regulatory mechanisms solely from transcriptional output (**Fig 1C**) [31–34]. This model uses two parameters to represent TF effects on transcription initiation: *β* and *α* for modulating RNAP–promoter binding and promoter escape, respectively. By applying GM to analyze <100 promoter variants regulated by each of 11 TFs in *Escherichia coli*, they reported that both repressor- and activator-regulated promoters exhibit an inverse scaling relationship between basal promoter strength and regulatory fold change (defined as the ratio of regulated to basal expression) [34]. Contrary to the widely accepted physical-blocking repression mechanisms, GM suggests repressors reduce transcription by trapping RNAP at the promoter through overstabilization of the RNAP–promoter complex.

Although GM bypasses the need for prior knowledge of TF function, its mechanistic interpretations of TF regulation contradict the consensus established through extensive biochemical characterization [35–41]. Additionally, our recent sort-seq characterization of >16,000 promoter variants per TF [30], compared with <100 in the GM study [34], revealed an intermediate optimum between basal promoter strength and fold change (termed peaked tradeoff), inconsistent with the GM-predicted universal inverse scaling trend. To examine the generality of this prediction and the credibility of the GM-inferred RNAP-trapping repression mechanism, here we compile and analyze 30 TF-regulated promoter datasets from four prior studies [30,34,42,43]. We then fit these datasets with GM and the conventional RM and AM and derive a mathematical solution linking basal expression to fold change. We find that over one third of them exhibit a peaked tradeoff between basal promoter strength and fold change. We further identify the assumption that constrains GM to the inverse scaling trend and show that its promoter-escape parameter *α* can be removed without compromising model-fitting performance. Together, these results establish basal promoter strength as a key design parameter for transcriptional regulation and highlight the importance of data coverage and model formulation in obtaining reliable model inferences.

## Methods

### Compilation of promoter datasets

Each of the 30 datasets (termed DS01–30; **Table 1**) derives from a single TF-regulated promoter library characterized in *E. coli*, except DS27 [30,34,42,43]. Variants within a library share a fixed promoter context and differ in the sequence composition of the −35 and/or −10 elements. DS27 pools two small libraries with the same promoter context but distinct Crp TFBS positions downstream of the transcription start site (+1). Expression from each promoter variant was quantified by sort-seq or fluorescence- or enzyme-based reporter assays under two conditions: (1) basal expression (*E*_TF–_), in which TF–promoter binding is prevented by TF gene knockout or addition of a ligand; and (2) regulated expression (*E*_TF+_), in which the TF can bind and regulate the promoter, with or without a ligand. Measurements were reported in arbitrary units different across datasets, with sample sizes ranging from 21 to 16,380. Among the datasets, each of the two subsets, DS08–DS13 and DS18–DS21, derives from a distinct promoter library assayed in different growth media; the promoter contexts for the remaining datasets are all unique, although some of them are regulated by the same TF.

**Table 1.** TF-regulated promoter datasets from prior studies.

| Dataset | TF | Regulation | Trend<sup>a</sup> | Growth medium<sup>b</sup> | Measurement | Sample size | Mutation |  | Source |
| --- | --- | --- | --- | --- | --- | --- | --- | --- | --- |
|  |  |  |  |  |  |  | -35 | -10 |  |
| DS01 | TetR | Repression | PT | LB+Glu | Sort-seq | 16,380 | + | + | [30] |
| DS02 | LuxR | Activation | PT | LB+Glu | Sort-seq | 16,363 | + | + | [30] |
| DS03 | CueR | Activation | PT | LB+Glu | Sort-seq | 16,368 | + | + | [30] |
| DS04 | AcrR | Repression | IS | M9+Glu | Fluorescent reporter | 49 | + | - | [34] |
| DS05 | AgaR | Repression | IS | M9+Glu | Fluorescent reporter | 42 | + | - | [34] |
| DS06 | AscG | Repression | PT | M9+Glu | Fluorescent reporter | 78 | + | - | [34] |
| DS07 | GntR | Repression | IS | M9+Glu | Fluorescent reporter | 89 | + | - | [34] |
| DS08 | LacI | Repression | PT | M9+Ace | Fluorescent reporter | 84 | + | - | [34] |
| DS09 | LacI | Repression | IS | M9+Ara | Fluorescent reporter | 85 | + | - | [34] |
| DS10 | LacI | Repression | IS | M9+Gal | Fluorescent reporter | 87 | + | - | [34] |
| DS11 | LacI | Repression | IS | M9+Glu | Fluorescent reporter | 86 | + | - | [34] |
| DS12 | LacI | Repression | IS | M9+Gly | Fluorescent reporter | 86 | + | - | [34] |
| DS13 | LacI | Repression | IS | M9+Pyr | Fluorescent reporter | 86 | + | - | [34] |
| DS14 | MngR<sup>c</sup> | Repression | IS | M9+Glu | Fluorescent reporter | 53 | + | - | [34] |
| DS15 | MngR<sup>d</sup> | Repression | IS | M9+Glu | Fluorescent reporter | 74 | + | - | [34] |
| DS16 | PdhR | Repression | IS | M9+Glu | Fluorescent reporter | 54 | + | - | [34] |
| DS17 | UlaR | Repression | IS | M9+Glu | Fluorescent reporter | 41 | + | - | [34] |
| DS18 | CpxR | Activation | IS | M9+Ara | Fluorescent reporter | 91 | + | - | [34] |
| DS19 | CpxR | Activation | IS | M9+Gal | Fluorescent reporter | 87 | + | - | [34] |
| DS20 | CpxR | Activation | IS | M9+Glu | Fluorescent reporter | 91 | + | - | [34] |
| DS21 | CpxR | Activation | IS | M9+Gly | Fluorescent reporter | 91 | + | - | [34] |
| DS22 | MetR | Activation | IS | M9+Glu | Fluorescent reporter | 58 | + | - | [34] |
| DS23 | SoxS<sup>c</sup> | Activation | PT | M9+Glu | Fluorescent reporter | 41 | + | - | [34] |
| DS24 | SoxS<sup>d</sup> | Activation | IS | M9+Glu | Fluorescent reporter | 43 | + | - | [34] |
| DS25 | AraC | Activation | PT | M9+Gly+CAA | Fluorescent reporter | 48 | + | + | [43] |
| DS26 | LasR | Activation | PT | M9+Gly+CAA | Fluorescent reporter | 48 | + | + | [43] |
| DS27 | Crp<br>(+1.5/+5.5)<sup>e</sup> | Repression | IS | RDM+Glu | β-galactosidase | 39 | + | - | [42] |
| DS28 | Crp<br>(-61.5)<sup>e</sup> | Activation | PT | RDM+Glu | β-galactosidase | 44 | + | + | [42] |
| DS29 | Crp<br>(-71.5)<sup>e</sup> | Activation | PT | RDM+Glu | β-galactosidase | 35 | + | + | [42] |
| DS30 | Crp<br>(-41.5)<sup>e</sup> | Activation | PT | RDM+Glu | β-galactosidase | 21 | - | + | [42] |

<sup>a</sup>PT, peaked tradeoff; IS, inverse scaling.
<sup>b</sup>Glu, glucose; Ace, acetate; Ara, arabinose; Gal, galactose; Gly, glycerol; Pyr, pyruvate; CAA, casamino acids; LB, lysogeny broth; M9, M9 minimal medium; RDM, rich defined medium.
<sup>c</sup>The promoter context is the synthetic promoter DL5p.
<sup>d</sup>The promoter context is the native promoter MngRp (DS15) or FldAp (DS24).
<sup>e</sup>Parenthetical values indicate the center position of the Crp TFBS relative to the transcription start site (+1), with the two values for DS27 corresponding to two pooled libraries that differ in TFBS position.

### Model formulation

We reconstructed RM, AM, and GM based on the formulation described in the source studies and fitted them to the compiled promoter datasets (**Table 1**). The three models share a theoretical framework which defines four promoter states: unbound, RNAP-bound, TF-bound, and co-bound (**Fig 1**) [11]. AM and GM have all four states whereas RM excludes the co-bound state because it assumes that RNAP and repressor binding to the promoter are mutually exclusive (**Fig 1A**). In these models, each promoter state (*i*) has a transcription rate *r*_i_ and a Boltzmann weight *P*_i_ proportional to its equilibrium probability. The Boltzmann weights of the unbound, RNAP-bound, TF-bound, and co-bound states are defined as 1, *P*_P_, and *P*_T_, and *β*·*P*_P_·*P*_T_, respectively. Here, *P*_P_ = [RNAP]/*K*_P_, and *P*_T_ = [TF]/*K*_T_, in which [RNAP] and [TF] are the concentrations of RNAP and TF competent for promoter binding, respectively, and *K*_P_ and *K*_T_ are the dissociation constants for their corresponding DNA binding sites. The parameter *β* describes TF influence on RNAP–promoter binding in the co-bound state. Regarding *r*_i_, RM and AM assign a transcription rate *r*_0_ to the unbound and TF-bound states, accounting for background transcription beyond the defined promoter region [21,42], whereas the transcription rate is *r*_max_ in the RNAP-bound and co-bound states, with *r*_0_ ≪ *r*_max_. GM sets *r*_0_ = 0 because promoter strength measurements in its associated datasets were obtained after subtraction of *E. coli* cellular autofluorescence (**Table 1**) [34]. GM further assigns a parameter *α* to describe TF modulation of promoter escape, which alters the transcription rate in the co-bound state (i.e., *α*·*r*_max_) [31–34]. Of AM and GM, AM assumes that all activators stabilize RNAP– promoter binding (*β* > 1), whereas GM uses both *β* and *α* to describe transcriptional regulation by repressors and activators with the following mechanistic interpretations: *β* > 1 and *β* < 1 for stabilization and destabilization of RNAP–promoter binding, respectively; *α* > 1 and *α* < 1 for acceleration and deceleration of promoter escape, respectively.

Based on the formulation described above, the magnitudes of *E*_TF–_ and *E*_TF+_ in each model are computed as probability-weighted sums of transcription rates across promoter states accessible under each condition (**Fig 1**), with the general form:

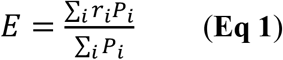

### Model fitting

Model fitting was carried out independently for each dataset, following the procedure of Forcier et al. [42]. Promoter strength measurements were log_10_-transformed before fitting. RM and AM were fitted to repressor-regulated (**Fig 2A**) and activator-regulated datasets (**Fig 2B**), respectively, whereas GM was fitted to both sets (**Fig 2**). *P*_P_ was fitted for each promoter variant because its value varies with promoter sequences, whereas the remaining free model parameters were fitted as single values shared across variants. All parameters were constrained to non-negative values and estimated by minimizing the sum of mean squared errors for *E*_TF−_ and *E*_TF+_, using gradient descent with PyTorch’s Adam optimizer. Model-fitting performance was quantified by the coefficient of determination (*R*^2^), root mean squared error (RMSE), and mean absolute error (MAE). These performance metrics and model-fitted values of parameters *α* and *β* are reported in **Fig 3** and **Fig 4**, respectively.

**Fig 2.**
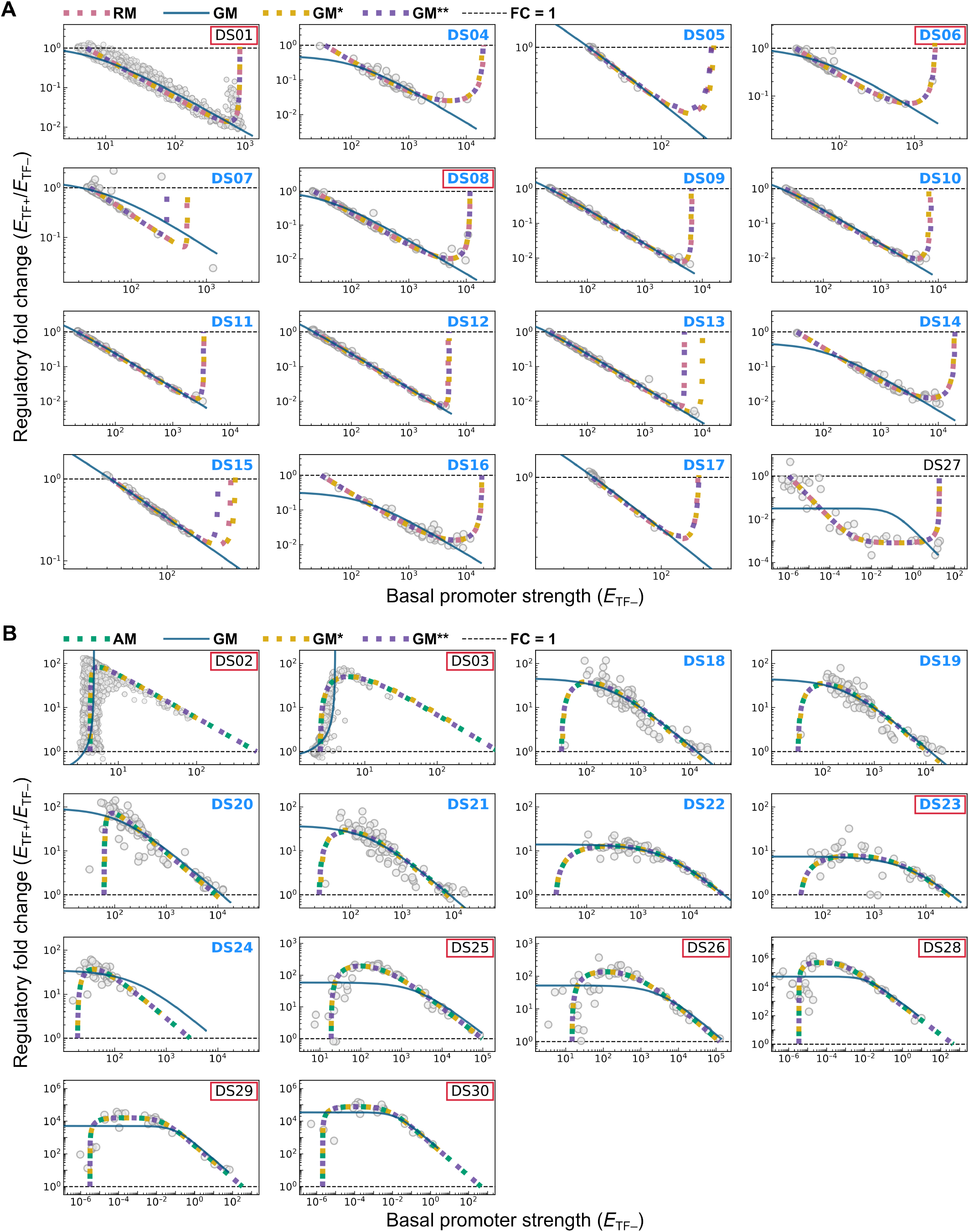
Modeling the relationship between basal promoter strength and regulatory fold change. **(A)** Model fitting of RM, GM, GM*, and GM** to 16 repressor-regulated datasets. **(B)** Model fitting of AM, GM, GM*, and GM** to 14 activator-regulated datasets. In **(A)** and **(B)**, experimental data are indicated by circles, regulatory fold change (*FC*) = 1 is marked by a dashed line, and model-fitting curves are shown as a solid line (GM) or colored dotted lines (RM, AM, GM*, and GM**). Datasets from the GM study are indicated by blue texts [34]. Datasets showing a peaked-tradeoff relationship are indicated by red boxes.

**Fig 3.**
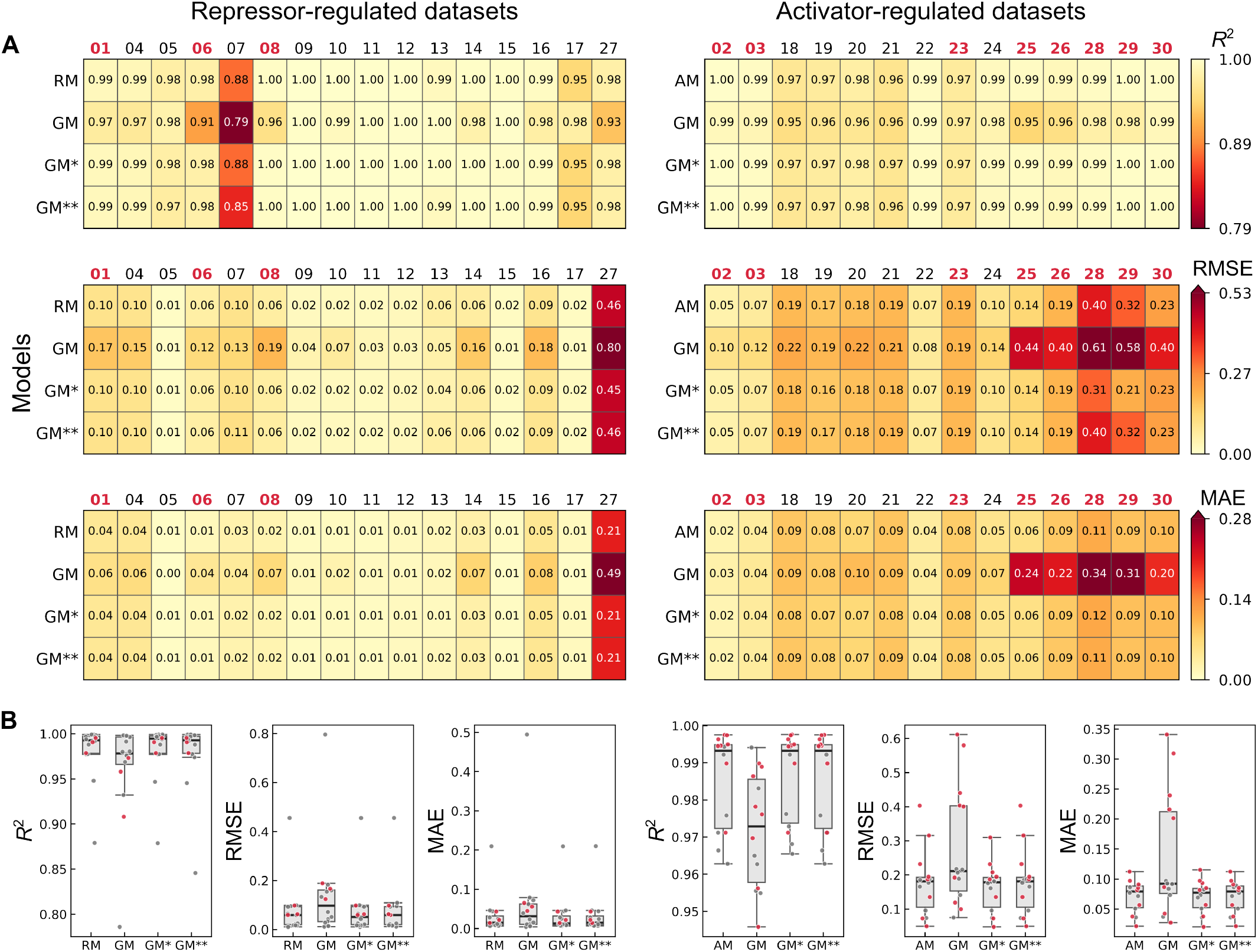
Model performance. **(A)** Model performance for each dataset, quantified by *R*^2^, RMSE, and MAE. Datasets showing a peaked-tradeoff relationship are indicated by red texts. **(B)** Distribution of each performance metric across datasets. Dots, boxes, whiskers, and central bars represent individual datasets, interquartile ranges, 1.5-fold interquartile ranges, and medians, respectively. Datasets showing a peaked-tradeoff relationship are indicated by red dots. In **(A)** and **(B)**, the left half shows RM, GM, GM*, and GM** fitted to repressor-regulated datasets, and the right half shows these models fitted to activator-regulated datasets.

**Fig 4.**
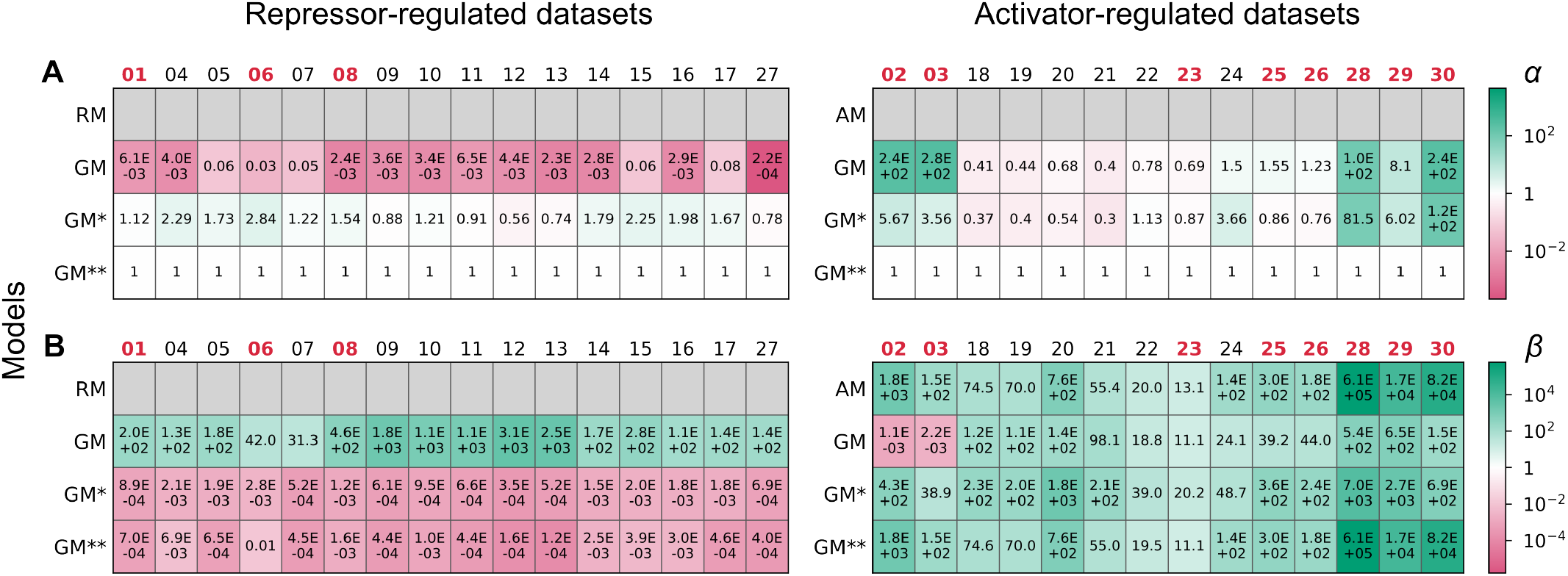
Model-estimated TF parameters. Estimates of parameters *α* **(A)** and *β* **(B)** represent TF influences on promoter escape and RNAP– promoter binding, respectively. In **(A)** and **(B)**, the left half shows RM, GM, GM*, and GM** fitted to repressor-regulated datasets, and the right half shows these models fitted to activator-regulated datasets. Datasets showing a peaked-tradeoff relationship are indicated by red texts. Gray cells denote parameters not included in the corresponding models, and *α* is fixed at 1 in GM**.

## Results and Discussion

### Intermediate optima between basal expression and fold change occur in over one third of datasets

We directly examined the relationship between basal expression (*E*_TF–_) and regulatory fold change (*FC* = *E*_TF+_/*E*_TF–_) for each promoter variant across the 30 datasets (represented by circles in **Fig 2**). Among repressor-regulated datasets (**Fig 2A**), *FC* declines as *E*_TF–_ increases in 13 of 16 cases (**Table 1**), matching the inverse relationship reported in the GM study [34]; yet the remaining datasets (DS01, DS06, and DS08) exhibit a clear intermediate optimum (minimum *FC* for repression), consistent with the peaked tradeoff observed in our prior work [30]. Intermediate optima (maximum *FC* for activation) is more common among activator-regulated datasets, occurring in 8 of 14 cases (**Table 1**; **Fig 2B**). Apparent overrepresentation of inverse scaling among repressor-regulated relative to activator-regulated datasets may be attributed to data sources and promoter library designs. The GM study [34] contributes to 21 datasets: 14 are repressor datasets (DS04–DS17) with 12 exhibiting inverse scaling, and 7 are activator datasets (DS18–DS24) with 6 exhibiting inverse scaling. Contrary to this predominance of inverse scaling, 8 of 9 datasets from the other studies show peaked tradeoff [30,42,43]. The sole exception shares a common feature with those from the GM study in that their promoter libraries bear sequence variation only in the –35 element. Previous experimental and modeling studies indicate greater importance of –10 than –35 elements in RNAP–promoter interactions [30,44–46]. Accordingly, promoter libraries generated by mutagenesis of just the –35 element may sample a narrower functional range, revealing only the inverse portion rather than the full peaked-tradeoff relationship between *E*_TF–_ and *FC*.

Collectively, 11 of the 30 promoter datasets exhibit peaked tradeoff, challenging the universality of the inverse scaling trend proposed in the GM study [34]. To test our narrow-sampling hypothesis and determine why GM predicted the inverse trend, we next fitted RM, AM, and GM to these datasets and examined potential issues in GM formulation.

### Conventional models outperform GM despite having fewer parameters

We fitted RM and GM to repressor datasets (**Fig 2A**), fitted AM and GM to activator datasets (**Fig 2B**), and examined the agreement between model fits and data distributions. Across repressor datasets, RM fits overall outperform GM fits in terms of *R*^2^ (0.88–1.00 vs 0.79–1.00), RMSE (0.01–0.46 vs 0.01–0.80), and MAE (0.01–0.21 vs 0.00–0.49) (**Fig 3A** and **3B**). RM-fitted curves are concave up, predicting an intermediate optimum between *E*_TF–_ and *FC*, while GM-fitted curves decline monotonically, predicting an inverse scaling relationship (**Fig 2A**). In parallel, across activator datasets, AM fits overall outperform GM fits in terms of *R*^2^ (0.97–1.00 vs 0.95–0.99), RMSE (0.05–0.40 vs 0.08–0.61), and MAE (0.02–0.11 vs 0.03–0.34) (**Fig 3A** and **3B**). AM-fitted curves are concave down, predicting an intermediate optimum between *E*_TF–_ and *FC*, while most GM-fitted curves decline monotonically, predicting an inverse scaling relationship (**Fig 2B**), with the exception of DS02 and DS03 in which GM-fitted curves unexpectedly rise monotonically. Relative to the RM–GM difference in fitting performance for repressor datasets, the AM– GM difference is more substantial (**Fig 3B**), reflecting the higher prevalence of peaked tradeoff among activator datasets (8/14) than repressor datasets (3/16) (**Fig 2** and **Table 1**). Taken together, the conventional models, RM and AM, outperform GM in fitting performance despite having fewer parameters.

### Sampled promoter function range shapes observed data distributions

Since the conventional models fit the empirical data better than GM, we used *P*_p_, the Boltzmann weight of the RNAP-bound state inferred by RM and AM, as a measure of promoter function and examined its range sampled by promoter variants in each dataset. We chose model-inferred *P*_p_ rather than experimentally determined promoter strength (*E*_TF–_ and *E*_TF+_) because the latter poorly distinguishes variants whose *P*_p_ values differ but lie in the flat regions of the sigmoidal *P*_p_–*E*_TF–_ and *P*_p_–*E*_TF+_ curves predicted by RM and AM (**Fig 5**). RM predicts the same shape for the *P*_p_–*E*_TF–_ and *P*_p_–*E*_TF+_ curves, with the latter left-shifted along the log-scale *P*_P_ axis, whereas AM also predicts identical shapes for the two curves but with the *P*_p_–*E*_TF+_ curve right-shifted. This shift, reflecting TF regulation, is quantified by the regulation factor (*F*_reg_) derived by Bintu et al. for RM and AM, respectively [11,17]:

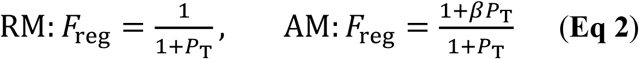

**Fig 5.**
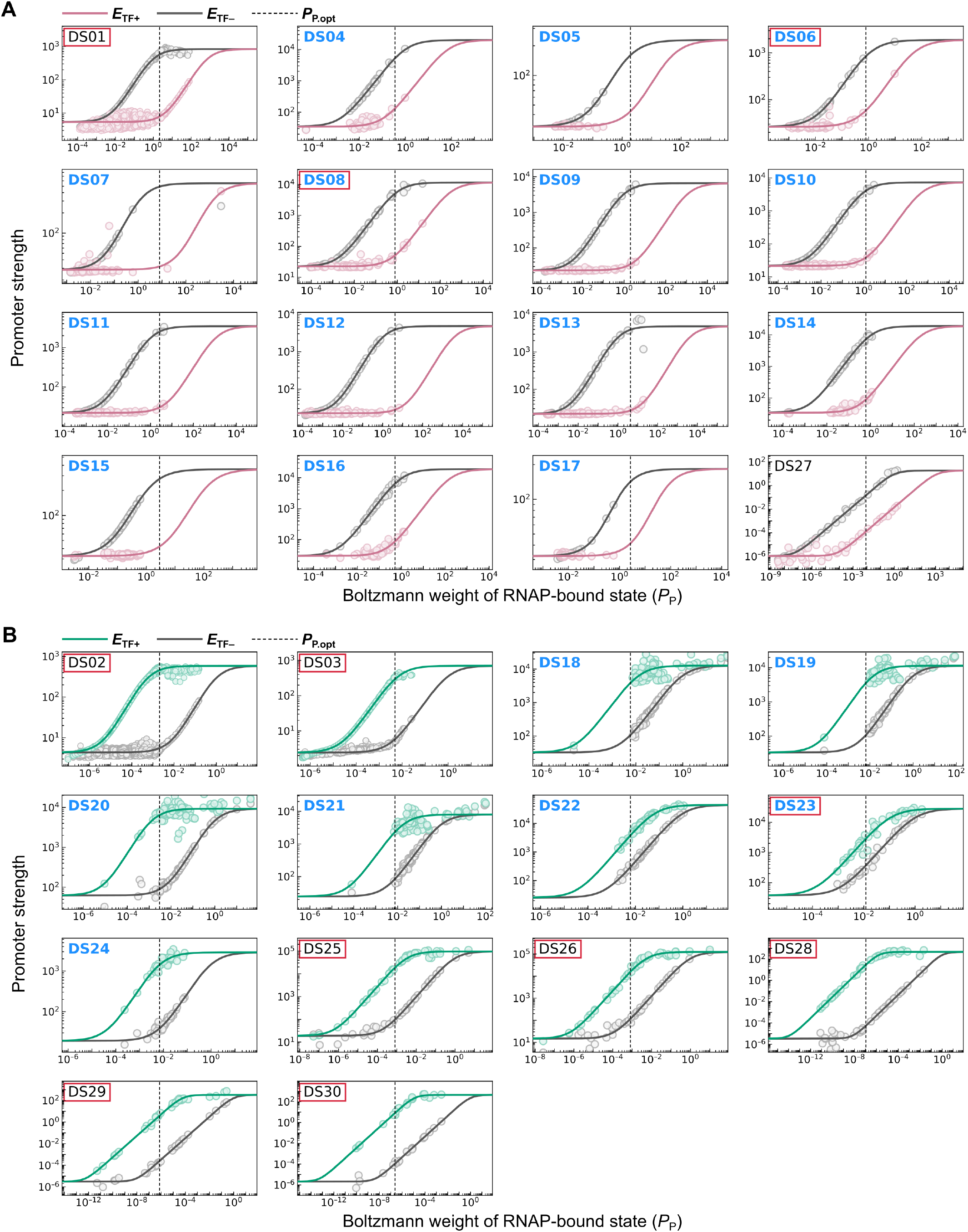
Relationship between RNAP-bound-state Boltzmann weight and promoter strength. **(A)** Model fitting of RM to 16 repressor-regulated datasets. **(B)** Model fitting of AM to 14 activator-regulated datasets. In **(A)** and **(B)**, circles indicate experimentally quantified basal (*E*_TF−_, gray) and regulated (*E*_TF+_, red and green, respectively) expression of individual promoter variants, plotted against model-inferred *P*_P_. Model-fitting curves are labeled by the corresponding colors. Vertical dashed lines mark *P*_P,opt_. Datasets showing a peaked-tradeoff relationship are indicated by red boxes.

Using *F*_reg_, *E*_TF–_ and *E*_TF+_ predicted by RM and AM can be reformulated in a unified form as:

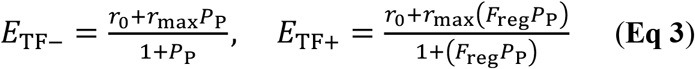

Furthermore, RM and AM predict a peaked-tradeoff relationship between *P*_p_ and *FC*, with concave-up and concave-down curve shapes, respectively (**Fig 6**). The *P*_p_ values (termed *P*_P.opt_) that yield the minimum *FC* in RM and the maximum *FC* in AM are derived in **S1 Text** section I as:

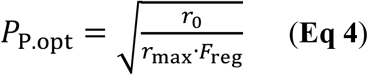

**Fig 6.**
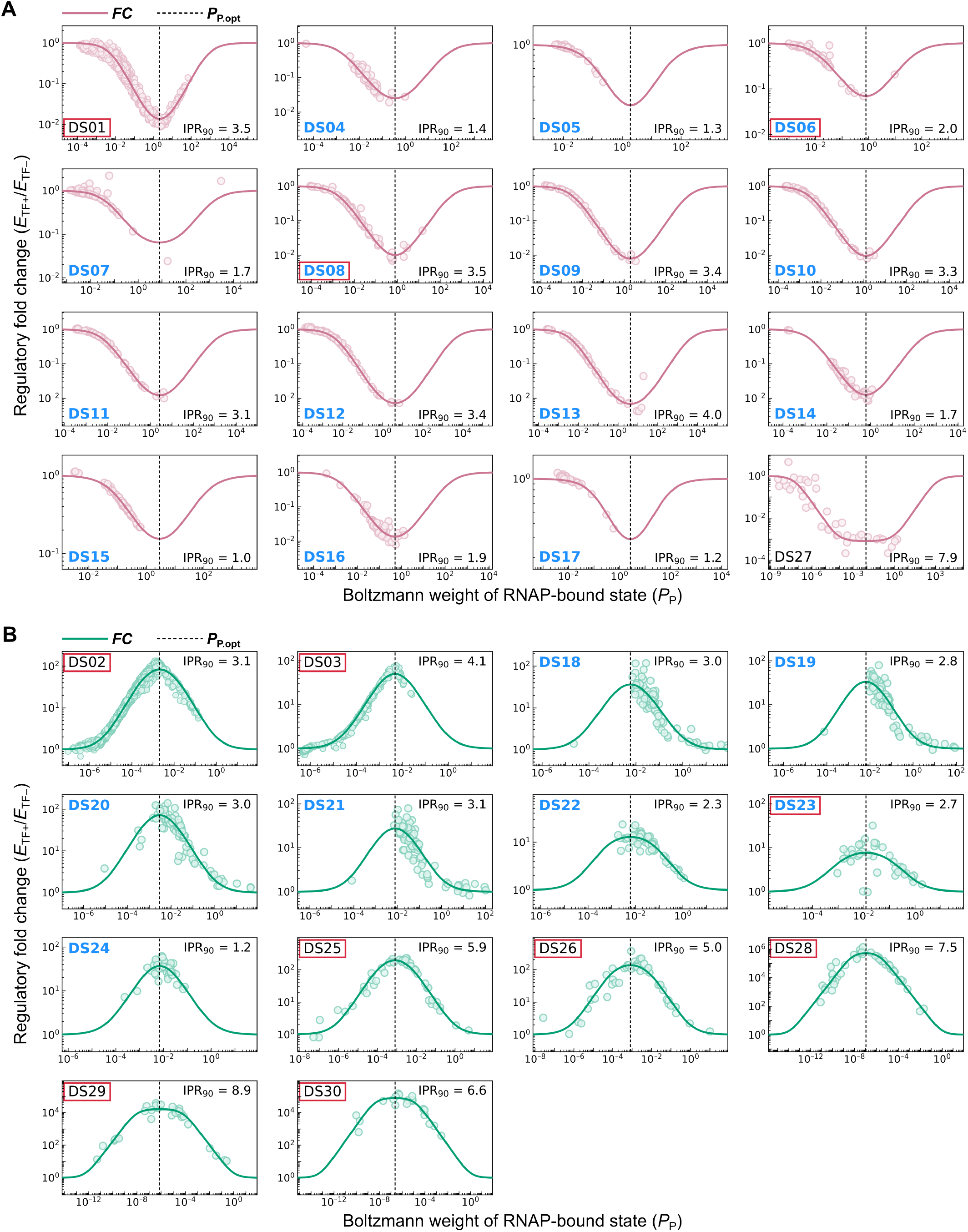
Relationship between RNAP-bound-state Boltzmann weight and regulatory fold change. **(A)** Model fitting of RM to 16 repressor-regulated datasets. **(B)** Model fitting of AM to 14 activator-regulated datasets. In **(A)** and **(B)**, circles indicate experimentally quantified regulatory fold change (*FC*) of individual promoter variants plotted against model-inferred *P*_P_. Model-fitting curves are shown as solid lines. Vertical dashed lines indicate *P*_P.opt_. Datasets from the GM study are indicated by blue texts [34]. Datasets showing a peaked-tradeoff relationship are indicated by red boxes. IPR_90_, 5th–95th percentile range of log_10_(*P*_P_) spanned by promoter variants within a dataset.

Notably, *P*_P.opt_ is located at the slope–plateau and bottom–slope junctions of the sigmoidal *P*_p_–*E*_TF–_ curve in RM and AM, respectively (**Fig 5**). TF-regulated promoters with *P*_P.opt_ provide the greatest response to transcriptional regulation, resulting in the maximum difference between *E*_TF–_ and *E*_TF+_. Together, these predictions reveal a general design principle of TF-regulated promoters, consistent with the conclusion of our previous study [30]. Additionally, for a dataset to reveal the peaked-tradeoff relationship between *E*_TF–_ and *FC*, the *P*_p_ values of sampled promoter variants have to span a range sufficient to cover *P*_P.opt_ and both sides of it (**Fig 6**). Datasets satisfying this requirement generally show such a peaked tradeoff in **Fig 2**. *P*_p_-based promoter analysis explains why datasets from the GM study mostly exhibit an inverse *E*_TF–_–*FC* relationship (18/21; **Fig 2**) [34]—although sampled promoter variants span a broad *E*_TF–_ range, they mostly cover the slope of the sigmoidal *P*_p_–*E*_TF–_ curve and fail to capture *P*_P.opt_ and both sides of it (**Fig 5** and **Fig 6**). The narrower data coverage of this study is also reflected in the 5th–95th percentile range of *P*_p_ (IPR_90_ = 1.0–4.0), compared with 3.1–8.9 for datasets from the remaining studies. In sum, the analyses above stress the need for broad data coverage to prevent a “blind men and the elephant” interpretation and establish a design principle that quantitatively connects basal promoter strength to transcriptional regulation.

### Relaxing a parameter constraint in GM changes its mechanistic interpretation

GM fits the relationship between *E*_TF–_ and *FC* less accurately than the conventional models because its fitted curves are monotonic (**Fig 2** and **Fig 3**), reflecting the constraint imposed by fixing parameter *r*_0_, which accounts for background transcription beyond the defined promoter region, at zero (**Fig 1**) [21,42]. Under the constraint of *r*_0_ = 0, mathematical derivation proves that the GM-predicted *E*_TF–_–*FC* relationship is monotonic, with its direction dependent solely on *β*: negative when *β* > 1 and positive when *β* < 1 (**S1 Text** section II). This derivation explains the unexpected positive monotonic *E*_TF–_–*FC* relationship predicted by GM for DS02 and DS03, in contrast with the negative monotonic trend for the remaining datasets.

To assess the impact of this model assumption, we fitted GM with unconstrained *r*_0_ (termed GM*) to the 30 datasets. GM*-fitted curves for repressor and activator datasets are concave up and down, respectively, closely matching the shapes of RM- and AM-fitted curves (**Fig 2**). In terms of fitting performance, GM* is comparable to RM and AM and better than GM for both repressor (*R*^2^: 0.88–1.00, RMSE: 0.01–0.45, MAE: 0.01–0.21) and activator datasets (*R*^2^: 0.97–1.00, RMSE: 0.05–0.31, MAE: 0.02–0.12) (**Fig 3**). These results indicate that the GM-predicted inverse scaling relationship between *E*_TF–_ and *FC* is an artifact of fixing *r*_0_ = 0 rather than a reflection of data properties. Although fixing *r*_0_ = 0 was justified in the GM study by fitting GM to datasets with cellular autofluorescence subtracted [34], this procedure does not eliminate residual transcription contributed by shadow promoters beyond the designated promoter region. Presence of background transcription activity is supported by the convergence of *FC* toward 1 at low basal promoter strength in their and others’ datasets (**Fig 2**), as this pattern suggests that transcription outside the TF-regulated promoters becomes dominant when expression from the designated promoters is weak.

In addition to reducing GM fitting performance, fixing *r*_0_ = 0 also alters model estimates of the TF parameters *α* (modulating promoter escape) and *β* (affecting RNAP–promoter binding), particularly for repressor datasets (**Fig 4**). In all 16 cases, GM estimates their *α* ≪ 1 and *β* ≫ 1, consistent with the trends reported in the GM study [34], whereas GM*, which outperforms GM, estimates *α* ≈ 1 and *β* ≪ 1. The GM study interpreted estimates of *β* ≫ 1 as strong support for an unconventional mechanism in which repressors inhibit transcription by overstabilizing RNAP–promoter binding in the co-bound state rather than by well-established physical-blocking mechanisms [2,47]. However, the contrasting fitting results between GM and GM*, which differ in just one model assumption, suggest that the proposed unconventional repression mechanism warrants careful reassessment. Moreover, GM* estimates *α* ≈ 1 for all repressor datasets, suggesting that this parameter, present in GM but absent from the conventional RM and AM, may be redundant. Given that the GM formulation is identical to AM except for including *α* and the *r*_0_ = 0 constraint (**Fig 1**), we then removed both from GM and fitted the resulting model (termed GM**, equivalent to AM) to both repressor and activator datasets.

### The promoter-escape parameter is dispensable for GM performance and inference

GM**, with *α* abolished (fixing *α* = 1) and *r*_0_ unconstrained, outperforms GM in fitting performance across all datasets, with *R*^2^ of 0.85–1.00 vs 0.79–1.00, RMSE of 0.01–0.46 vs 0.01–0.80, and MAE of 0.01–0.21 vs 0.00–0.49 (**Fig 3**). GM** fitting performance is comparable to that of RM, AM, GM* (**Fig 3B**), and its fitted curves exhibit peaked-tradeoff shapes similar to those of these models for the corresponding datasets (**Fig 2**). Given identical model formulation, similar fits by GM** and AM are expected. Nevertheless, the similar fits of GM** and GM*, which differ only in the absence or presence of *α*, indicate that the model formulation from the GM study is unable to infer TF regulatory mechanisms from transcriptional output lacking kinetic information. Considered together, predictions by the conventional models and revised GM variants, along with empirical data distributions presented above, support a peaked-tradeoff rather than inverse scaling relationship between *E*_TF–_ and *FC*.

## Funding

This research is supported by Taiwan National Science and Technology Council grants (112-2311-B-002-011-MY3, 114-2311-B-002-002, 114-2811-B-002-189, 115-2311-B-002-001, 115-2311-B-002-011-MY3, 115-2811-B-002-025).

## Author Contributions

Conceptualization: H.H.D.C.; Methodology: S.T.A.K.; Investigation: S.T.A.K., C.P.H., H.H.D.C.; Formal analysis: S.T.A.K.; Visualization: S.T.A.K., H.H.D.C.; Supervision: H.H.D.C.; Funding acquisition: H.H.D.C.; Writing – Original Draft: S.T.A.K., H.H.D.C.; Writing – Review & Editing: S.T.A.K., H.H.D.C.

## Competing Interests

The authors declare no competing interests.

## S1 Text. Analytical properties of regulatory fold change

### Section I. The conventional models admit a closed-form solution for *P*_**P.opt**_

Under both RM and AM formulation, the basal (*E*_TF−_) and regulated (*E*_TF+_) promoter strengths are given by (**Eq 3** in the main text)

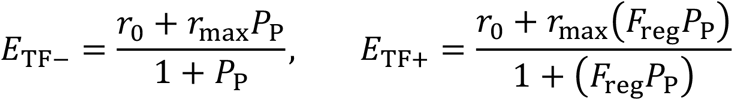

where *r*_0_, *r*_max_, and *F*_reg_ are constants within a dataset, whereas *P*_P_ varies across promoter variants. All of these quantities are positive, with *r*_0_ < *r*_max_, *F*_reg_ < 1 in the RM, and *F*_reg_ > 1 in the AM. A closed-form solution for *P*_P_ at which *FC* reaches its minimum in the RM and its maximum in the AM is derived below. To simplify the subsequent calculation, we let *μ* = *r*_0_/*r*_max_ (0 < *μ* < 1). The regulatory fold change is defined as *FC* = *E*_TF+_/*E*_TF−_, given by

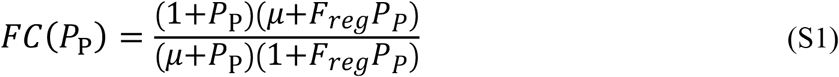

Differentiating ln(*FC*) is more tractable than differentiating *FC* directly, and the two derivatives are related by the chain rule,

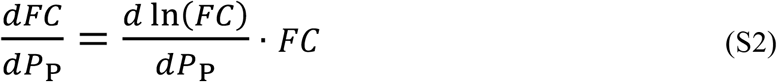

Since *FC* > 0, the two derivatives share the same sign,

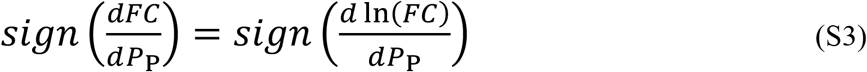

so the analysis below is carried out on ln(*FC*), which is given by

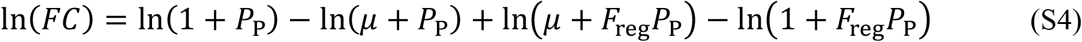

Differentiating Eq S4 with respect to *P*_P_ yields

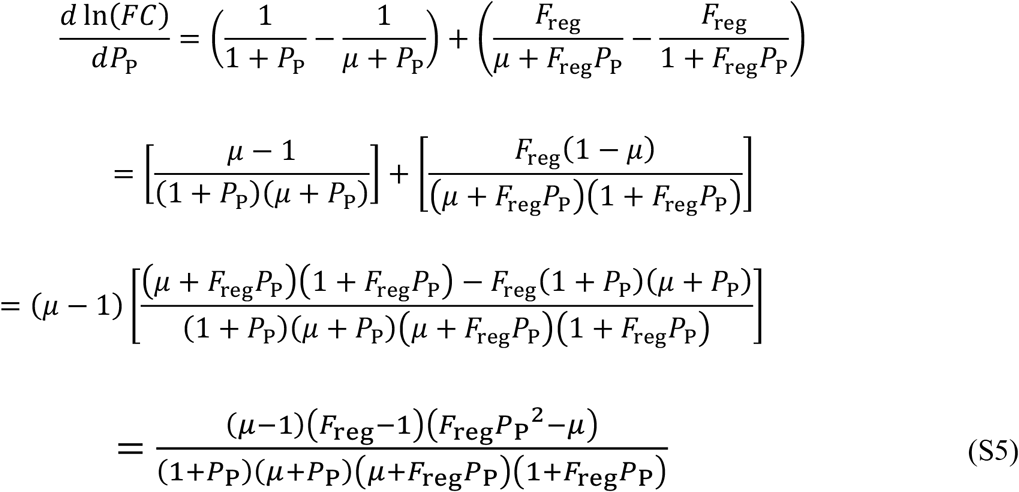

Substituting Eq S5 into Eq S3 and noting that the denominator is positive for all *P*_P_ > 0 whereas (*μ* − 1) is negative, the sign of *dFC*/*dP*_P_ is given by

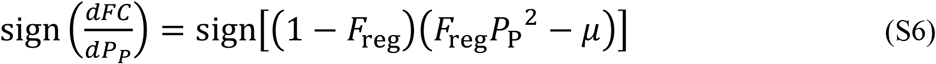

Because 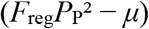 increases strictly with *P*_P_, it equals zero at a single positive value, denoted by *P*_P.opt_,

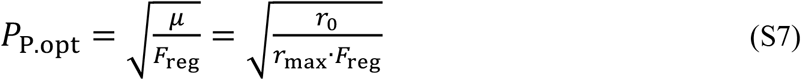

Under the RM model, (1 − *F*_reg_) is positive, whereas 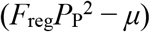 is negative for *P*_P_ < *P*_P.opt_ and positive for *P*_P_ > *P*_P.opt_. The derivative *dFC*/*dP*_P_ is therefore negative below *P*_P.opt_ and positive above it, making *P*_P.opt_ the unique global minimum of *FC*. By contrast, under the AM model, (1 − *F*_reg_) is negative, both signs are therefore reversed, and *P*_P.opt_ is the unique global maximum.

### Section II. Fixing *r*_0_ = 0 forces a monotonic *E*_TF−_–*FC* relationship in GM

Under the GM formulation, the basal (*E*_TF–_) and regulated (*E*_TF+_) promoter strengths are given by (**Fig 1C** in the main text)

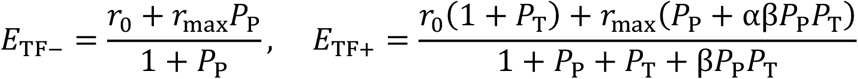

where *r*_0_, *r*_max_, *α, β*, and *P*_T_ are constant and *P*_P_ is variable within a dataset. The regulatory fold change is defined as *FC* = *E*_TF+_/*E*_TF−_. We show that the constraint *r*_0_ = 0 forces *FC* to be a monotonic function of *E*_TF−_ for any fixed set of {*α, β, P*_T_}. With *r*_0_ = 0 and *α, β, P*_T_ all positive, *FC* as a function of *P*_P_ is given by

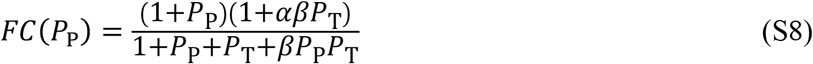

When *r*_0_ = 0, *P*_P_ = *E*_TF−_/(*r*_max_ – *E*_TF−_). Substituting this equation into Eq S8, *FC* as a function of *E*_TF−_ is given by

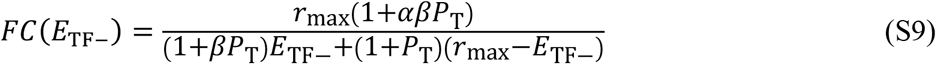

To simplify the subsequent calculation, we divide Eq S9 into a constant numerator and an *E*_TF−_-dependent denominator, letting *FC*(*E*_TF−_) = *c*/*f*(*E*_TF−_), where

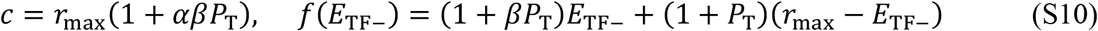

and differentiate *FC*(*E*_TF−_) with respect to *E*_TF−_ using the quotient rule,

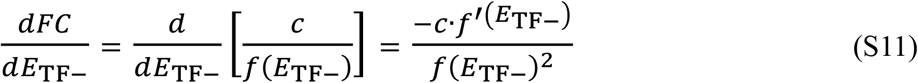

We first evaluate *f’*(*E*_TF−_) = (1+*βP*_T_) − (1+*P*_T_) = *P*_T_ (*β* − 1), so that

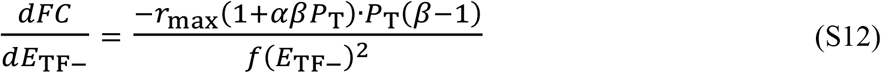

Because *f*(*E*_TF−_)^2^ > 0 and *r*_max_·(1+*αβP*_T_)·*P*_T_ > 0, the sign of *dFC*/*dE*_TF−_ is determined entirely by −(*β* − 1),

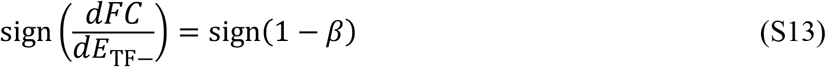

Since the sign of *dFC*/*dE*_TF−_ is independent of *E*_TF−_, it remains unchanged across all values of *E*_TF−_. Consequently, when *β* > 1, *FC* decreases monotonically with *E*_TF−_; when *β* < 1, *FC* increases monotonically with *E*_TF−_; and when *β* = 1, Eq S12 gives *dFC*/*dE*_TF−_ = 0, so *FC* remains constant with respect to *E*_TF−_. In every case, an interior extremum in *FC* as a function of *E*_TF−_ is mathematically precluded once *r*_0_ = 0 is imposed, regardless of the values of *α, β*, or *P*_T_.

## Notes

### Competing Interest Statement

The authors have declared no competing interest.

